# Horizontal gene transfers are widespread across termite genomes but do not confer metabolic innovations

**DOI:** 10.1101/2025.03.16.643567

**Authors:** Cong Liu, Simon Hellemans, Yukihiro Kinjo, Alina A. Mikhailova, Cédric Aumont, Yi Ming Weng, Aleš Buček, Filip Husnik, Jan Šobotník, Mark C. Harrison, Dino P. McMahon, Thomas Bourguignon

## Abstract

Horizontal gene transfer (HGT), the transmission of genetic material across species, is an important innovation source in prokaryotes. In contrast, its significance is unclear in many eukaryotes, including insects. Here, we used high-quality genomes of 45 termites and two cockroaches to investigate HGTs across blattodean genomes. We identified 289 genes and 2,494 pseudogenes classified into 168 orthologous groups originating from an estimated 281 HGT events. *Wolbachia* represented the primary HGT source, while termite gut bacteria and the cockroach endosymbiont *Blattabacterium* did not contribute meaningfully to HGTs. Most horizontally acquired genes descended from recent and species-specific HGTs, experienced frequent duplications and pseudogenizations, and accumulated substitutions faster than synonymous sites of native protein-coding genes. Genes frequently transferred horizontally to termite genomes included mobile genetic elements and genetic information processing genes. Our results indicate that termites continuously acquired genes through HGT, which they soon lost, leaving few horizontally acquired genes conserved across lineages.

## Introduction

Horizontal gene transfer (HGT) is the transmission of genetic material across species (Soucy et al. 2015; Tokuda and Shintani 2024). HGT has been playing an essential role in the evolution of bacteria and archaea, allowing for their rapid adaptation to new environments (Gogarten and Townsend 2005; Arnold et al. 2022). This is well illustrated by the largely HGT-driven spread of antibiotic resistance among human-associated microbes (Andam et al. 2011; Woods et al. 2020; McInnes et al. 2020). While the significant role of HGT in the evolution of prokaryotes has been recognized for many decades (Ochman et al. 2000; Koonin 2016), its importance and frequency in eukaryotes is less clear. The process of HGT also likely differs among lineages of eukaryotes, depending on various factors such as the presence of soma and germ line cells, the host environment, and the presence of intracellular heritable symbionts (Husnik and McCutcheon 2018; Keeling 2024).

The identification of HGTs in eukaryotes is often hampered by methodological artifacts, the presence of contaminants in draft genome assemblies, and the complex nature of eukaryotic genomes. Consequently, HGTs have been wrongly reported on multiple occasions (*e.g.*, Crisp et al. 2015; Salzberg 2017). Nevertheless, HGTs do occur in many eukaryotes, including insects, and can be confidently uncovered with appropriate methodologies (Nakabachi 2015; Husnik and McCutcheon 2018). For example, pea aphids encode carotenoid biosynthetic enzymes of fungal origin (Moran and Jarvik 2010), whiteflies possess a plant-derived gene participating in food detoxification (Xia et al. 2021), and lepidopteran genomes contain virus-derived genes providing protection against parasitoids (Gasmi et al. 2021). Insect genomes frequently contain sequences acquired from intracellular bacteria such as *Wolbachia*, which generally undergo rapid pseudogenization (Nakabachi 2015). Therefore, HGTs are known to occur in insects, but the overall dynamics of HGT, including the frequency, evolutionary significance, and fate of acquired genes, are largely unknown, primarily owing to the lack of comparative analyses among genomes of closely related species.

Termites (Insecta, Blattodea) are the second largest lineage of social insects and diverged from their sister group, the sub-social wood roach *Cryptocercus*, about 170 million years ago (Lo et al. 2000; Bourguignon et al. 2015). Termites and *Cryptocercus* live in symbiotic relationships with various organisms. They host gut symbionts that help digest plant materials, including bacteria, archaea, and protists present in all termites except the Termitidae (Brune and Dietrich 2015; Bignell 2016; Gile 2024). In addition, Macrotermitinae externally cultivate fungi, and their sister clade, Sphaerotermitinae, is hypothesized to have evolved an external symbiosis with bacteria (Bucek et al. 2019; Chouvenc et al. 2021). Termites are also frequently infected by *Wolbachia* (Bandi et al. 1997; Lo and Evans 2007; Salunke et al. 2010; Roy et al. 2015; Hellemans et al. 2019), an intracellular bacterium prevalent in terrestrial arthropods and nematodes (Kaur et al. 2021; Porter and Sullivan 2023). Finally, cockroaches and *Mastotermes* host *Blattabacterium*, an obligate nutrient-provisioning bacterial endosymbiont, which was lost in all non-*Mastotermes* termites (Sabree et al. 2009; Sabree et al. 2012; Neef et al. 2011; Kinjo et al. 2018). The long-term evolutionary associations between termites and various symbionts may provide opportunities for many HGTs into their genomes. Associations involving vertical transmission of gut microbes through the exchange of gut fluids between the colony founders and nestmates (Michaud et al. 2020; Sinotte et al. 2023; Arora et al. 2022, 2023; Gile 2024) and transovarial transmission in the case of *Wolbachia* and *Blattabacterium* could represent especially interesting sources of HGT, given the importance of these symbionts to termite physiology.

In this study, we systematically searched for horizontally acquired genes (including pseudogenes) in 45 high-quality genomes of termites representing the major termite lineages and two cockroach outgroups, *Cryptocercus meridianus* and *Blatta orientalis* (Liu et al. 2025) (Table S1). We screened the genomes with strict filtering criteria to prevent misidentifications caused by contaminations. We performed phylogenetic analyses to determine the origin of the identified horizontally acquired genes. Finally, we investigated the evolutionary fate of genes acquired by HGT.

## Results and discussion

### Robust detection of HAGPs

We used the alien index (AI) (Gladyshev et al. 2008; Yuan et al. 2023), which compares *e*-values of non-arthropod metazoan and non-metazoan homology hits, and the proportion of homology hits from non-arthropod metazoan and non-metazoan organisms to initially screen all genes annotated by Liu et al. (2025) across 45 termite and two cockroach genomes and identify potential HGTs. Genes suspected of originating from HGTs were validated using phylogenetic trees of their hierarchical orthologous groups (HOGs), delimited with OrthoFinder (Emms and Kelly 2019), and non-metazoan sequences. We identified 289 horizontally acquired genes and 2,494 horizontally acquired pseudogenes distributed across 1,313 contigs from 45 termite and two cockroach genomes (Tables S2-3). We confirmed the absence of false positives by inspecting the contigs containing horizontally acquired genes and pseudogenes (HAGPs) for potential contamination. Sequence properties of contigs with HAGPs corroborated that they are contamination-free portions of termite or cockroach nuclear genomes. First, all contigs containing HAGPs had a coding density lower than 70% in the annotation performed with the prokaryotic genome annotator PRODIGAL and 27% in the original genome annotation performed with the eukaryotic genome annotation pipeline of Liu et al. (2025) (Figure 1A), while the coding density of bacterial genomes is typically above 85% (Land et al. 2015). Second, HAGPs made up a maximum of 32.03% of contig length, and 99% of contigs containing HAGPs had more than two-thirds of their genes of native insect origin (Figure 1B). Third, all contigs containing genes and pseudogenes acquired by HGT also contained transposons in a proportion similar to that of cockroach and termite genomes suggesting a blattodean origin (Figure 1C). Fourth, less than 0.7% of the contig lengths aligned to *Wolbachia* genomes for 99% of the contigs containing HAGPs, and these proportions were consistently lower than 0.9% for *Blattabacterium* and termite gut bacteria (Figure 1D), indicating that the HGTs identified here were not contaminations from insect-associated symbionts. Fifth, the sequencing depth of contigs with HAGPs was generally within the range of genome sequencing depth: it varied between 0.62 and 2.08 times that of the genomes when the genome sequencing depth was above 30 (deep sequencing), and between 0.52 and 2.41 times when the sequencing depth was below 30 (shallow sequencing) (Figure 1E). Sixth, the sequencing coverage of contigs containing horizontally acquired genes or pseudogenes was high: in deep sequencing, reads mapped along more than 98.86% of the contig lengths for 90% of the contigs, while in shallow sequencing, reads mapped along more than 54.68% of the contig lengths for 90% of the contigs (Figure 1F). Seventh, the sequencing depth of HAGPs was generally similar to the genome sequencing depth, with 5% and 95% quantiles being 0.42 and 3.16 times that of the genomes in deep sequencing and 0 and 3.71 in shallow sequencing (Figure 1G). Eighth, HAGPs had high sequencing coverage: in deep sequencing, read coverage was 100% in more than 90% of the contigs, while in shallow sequencing, the coverage was above 39.45% in more than 90% of the contigs (Figure 1H). Overall, these results confirm that the HAGPs identified here are part of the termite and cockroach genomes.

**Figure 1.**
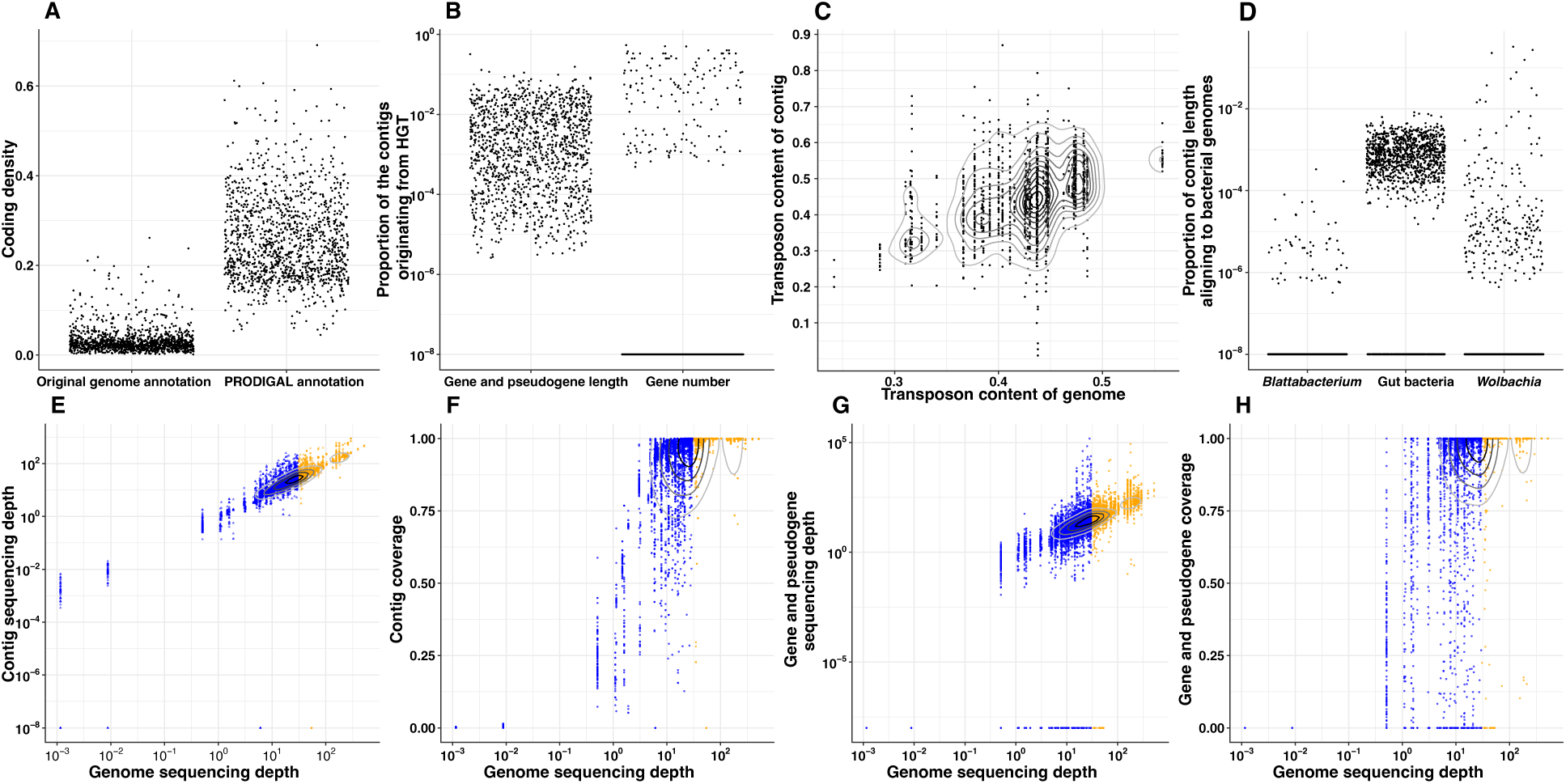
Validation of horizontal gene transfer events. Only contigs containing horizontally acquired genes or pseudogenes are shown. (A) Coding density of contigs estimated with the original genome annotations and the annotations obtained with PRODIGAL. (B) Proportion of contig length represented by HAGPs, and proportion of horizontally acquired genes. (C) Scatter density plot showing the transposon content of contigs and their corresponding genomes. (D) Proportion of contig length aligning to the genomes of *Wolbachia*, termite gut bacteria, and *Blattabacterium*. Scatter density plot showing: (E) the sequencing depth of contigs, (F) the coverage of contigs, (G) the sequencing depth of HAGPs, and (H) the coverage of HAGPs compared to their corresponding genomes. Blue dots indicate sequencing depth below 30 and orange dots sequencing depth above 30. Black counter lines indicate high density.

The 47 genomes analysed in this study were of variable quality (Liu et al. 2025). To assess the influence of genome quality on our ability to detect HGTs, we tested for a correlation between the number of HAGPs and four indexes of genome assembly and annotation quality reported by Liu et al. (2025) using Spearman’s rank correlation coefficient (*ρ*). There was no significant correlation between the number of horizontally acquired genes and pseudogenes and the N50 (P=0.501, *ρ* =0.101) or the L90 (P=0.357, *ρ* = 0.138) and a negative correlation with the BUSCO completeness (insecta_odb10) (Seppey et al. 2019) of genome assemblies (P=0.021, *ρ* = −0.337) and proteomes (P=0.002, *ρ* = −0.447). Therefore, genome quality mildly affected our ability to detect HGTs.

### Wolbachia is *the primary donor of horizontal gene transfers in termites*

The 289 horizontally acquired genes and 2,494 pseudogene copies found across our 47 genomes were clustered into 110 hierarchical orthologous groups (HOGs). Our phylogenetic analyses further split these HOGs into 168 sub-orthogroups forming sister clades to non-metazoan sequences, each representing at least one independent transfer to termite or cockroach genomes. Bacteria were the main contributor to HGTs as 125 sub-orthogroups were of bacterial origin, including 103 from *Wolbachia* and 17 that could not be assigned more precisely than to Pseudomonadota, the phylum to which *Wolbachia* belongs. Eight sub-orthogroups were of fungal origin, and 11 sub-orthogroups originated from plants. The remaining 24 sub-orthogroups originated from viruses.

*Wolbachia* is an intracellular bacterium that frequently infects arthropods (Kaur et al. 2021; Porter and Sullivan 2023), including termites (Bandi et al. 1997; Lo and Evans 2007; Salunke et al. 2010; Roy et al. 2015; Hellemans et al. 2019; Yashiro and Lo 2019). Six of the 47 genome assemblies used here also contained contigs with *Wolbachia* ribosomal RNA genes identified by Barrnap v.0.9 (Seeman 2013), which were excluded during HGT validation (Table S4), indicating *Wolbachia* infection. *Wolbachia* is transmitted vertically through the eggs (Kaur et al. 2021; Porter and Sullivan 2023), providing a potential route for HGTs into host genomes (Nakabachi 2015; Husnik and McCutcheon 2018; Keeling 2024). Furthermore, *Wolbachia* genomes are rich in mobile genetic elements that promote HGTs among strains (Ishmael et al. 2009; Scholz et al. 2020; Kaur et al. 2021), and possibly between *Wolbachia* and their hosts. The frequency of *Wolbachia* infection, their association with insect germline, and the abundance of mobile genetic elements potentially explain their predominant contribution to HGT events in termites, as is the case in other insects (Hotopp et al. 2007; Nikoh et al. 2008; Choi et al. 2015; Nakabachi 2015; Leclercq et al. 2016).

### No evidence of horizontal gene transfer from Blattabacterium and gut bacteria to termites

Surprisingly, while cockroaches and termites host bacterial symbionts vertically transmitted over tens of millions of years, including gut bacteria (Arora et al. 2022, 2023; Beránková et al. 2024) and *Blattabacterium* (Lo et al. 2003; Kinjo et al. 2021), these do not appear to contribute meaningfully to HGTs in termites. We found no evidence of their contribution to the pool of horizontally acquired genes identified in our 47 genomes. Furthermore, the inspection of local alignments of the genomes of 13 termite gut bacteria (Table S5) and 16 *Blattabacterium* strains (Table S6) to the 47 termite and cockroach genomes revealed no clear evidence of HGTs. Indeed, we found 350 termite and cockroach genes with coding regions partly aligning with termite gut bacterial genomes; however, homology searches against the non-redundant (nr) database indicated that the proportion of non-metazoan hits was systematically below 4%, except for two hits for which these proportions reached 8.8% and 22.6%. Similarly, there were three genes whose coding regions partly aligned with the genomes of *Blattabacterium*, with a proportion of non-metazoan homology hits lower than 0.2%. Furthermore, these alignments with bacterial genomes were short, as the length of alignments with identity above 20% was at most 417 and 1,180 bp for termite gut bacteria and *Blattabacterium* (Table S7), respectively. Overall, these results confirm that neither termite gut bacteria nor *Blattabacterium* significantly contributed to HGTs in termites.

The close association of *Blattabacterium* with cockroach eggs could theoretically promote HGTs to the host genomes, as is the case for *Wolbachia*. However, *Blattabacterium* and *Wolbachia* are different in two critical ways. First, *Blattabacterium* is confined to specialized host cells and their transmission is tightly regulated (Sacchi et al. 2000; Noda et al. 2020), whereas *Wolbachia* infects various tissues and switches hosts frequently (Kaur et al. 2021; Porter and Sullivan 2023). Second, *Blattabacterium* genomes are highly conserved and streamlined, containing no mobile elements or bacteriophages (Sabree et al. 2009; Patino-Navarrete et al. 2013; Kinjo et al. 2018), which likely hinders their insertion into the host genomes. In contrast, *Wolbachia* genomes vary in gene content (Ishmael et al. 2009; Scholz et al. 2020) and are rich in mobile genetic elements (Kaur et al. 2021; Porter and Sullivan 2023). The absence of gene transfer from *Blattabacterium* to the host genomes is reminiscent of other obligate endosymbionts, which also appear to rarely be the source of HGTs to the genomes of their hosts (Nikoh et al. 2010; Husnik et al. 2013; Luan et al. 2015; Sloan et al. 2014).

The absence of HGTs from termite gut bacteria despite their association with termites over geological timescales (Arora et al. 2022, 2023; Beránková et al. 2024) might be explained by the lack of close contact with host germline cells. The gut bacteria of termites and *Cryptocercus* are transmitted across host generations through the exchange of hindgut fluids (Michaud et al. 2020; Nalepa et al. 2001; Sinotte et al. 2023). Overall, our results highlight the absence of genetic material exchange between insects and their specialized mutualistic bacteria.

### Horizontally acquired genes and pseudogenes (HAGPs) are unevenly distributed among termites

The termite and cockroach genomes analyzed here contained between 0 to 53 horizontally acquired genes (mean: 6.15, median: 3) and 3 to 273 pseudogenes (mean: 53.06, median: 45) (Figures 2A-C). Five species (*Cryptotermes longicollis*, *Neotermes castaneus*, *Stylotermes halumicus*, *Coptotermes testaceus*, and *Heterotermes tenuis*) only had horizontally acquired pseudogenes but no genic copy. In contrast, the most extreme cases, *Leptomyxotermes doriae* and the cockroach *Blatta orientalis*, each contained 53 and 38 horizontally acquired genes (Figure 2B), and *Anoplotermes pacificus* and *Isognathotermes planifrons* contained 273 and 158 pseudogenes originating from HGTs, respectively (Figure 2C). The HAGPs of *L. doriae* belonged to 56 sub-orthogroups, the largest number found in a single termite genome (Figure 2D). Furthermore, the horizontally acquired sub-orthogroups were represented by multiple copies of genes and pseudogenes in many genomes (Figure 2E). The average copy number of genes and pseudogenes per sub-orthogroup per genome ranged from 1.20 to 14.84, indicating high rates of duplications following HGTs. The number of pseudogenized copies originating from HGTs outnumbered the number of horizontally acquired genes by a factor of ten, and more than half of the sub-orthogroups were only represented by pseudogenes in 40 genomes (Figure 2F), indicating that most genes of foreign origin face pseudogenization. Overall, the numbers of horizontally acquired genes found in termites are comparable with other insects (Li et al. 2022), and our results highlight the non-functionality and transient nature of most horizontally acquired genes.

**Figure 2.**
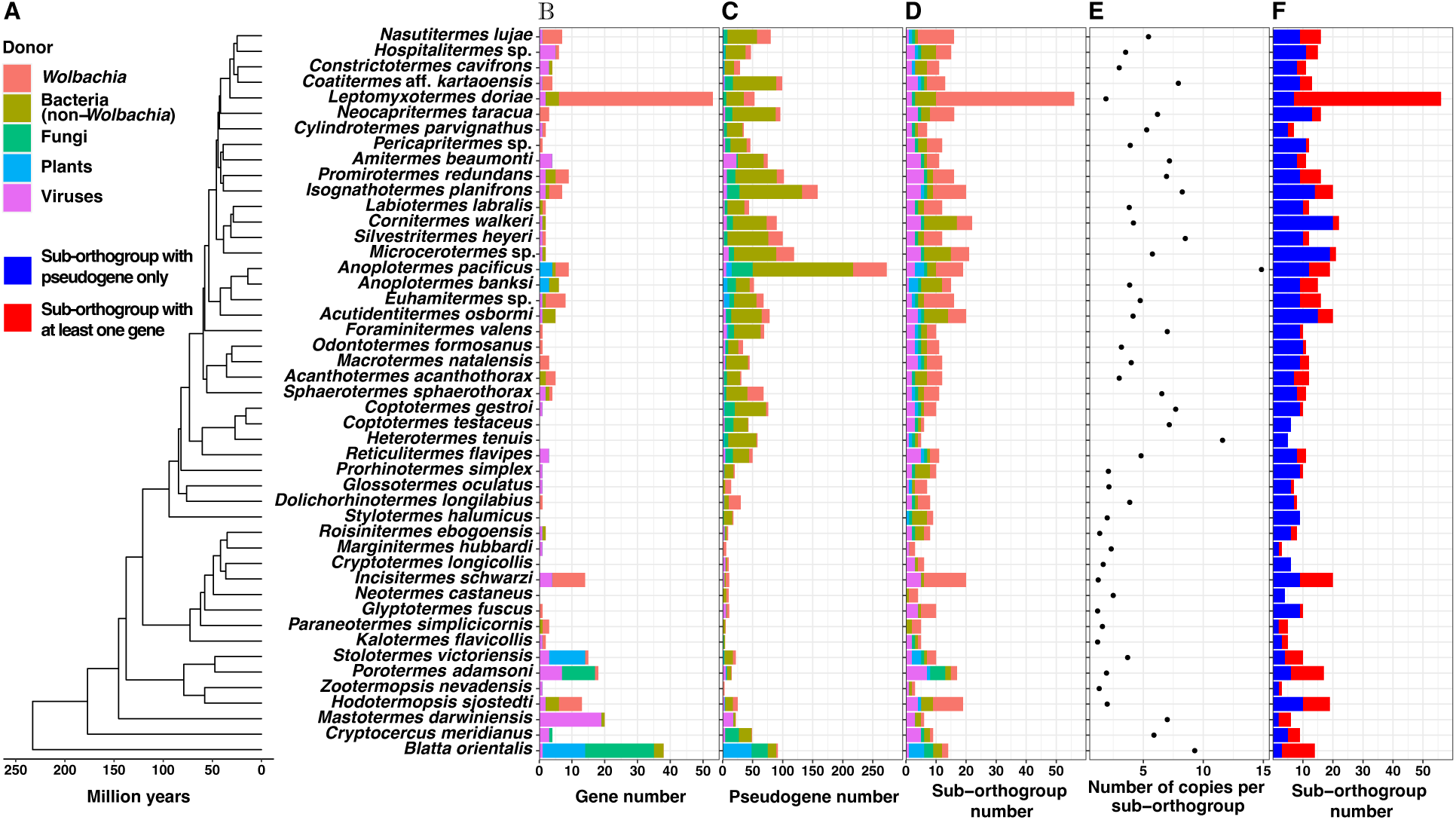
Frequency of HAGPs in termite genomes. (A) Time-calibrated species tree of the 45 termites and two cockroaches used in this study. Number of horizontally acquired (B) genes, (C) pseudogenes, and (D) sub-orthogroups. (E) Average copy number per sub-or-thogroup. (F) Number of sub-orthogroups only represented by pseudogenes (blue) and rep-resented by at least one gene (red).

### *Insertions of* Wolbachia *genome fragments in termite genomes*

The genomes of many insect species contain large insertions of *Wolbachia* genome fragments (Hotopp et al. 2007; Nikoh et al. 2008; Brelsfoard et al. 2014; Klasson et al. 2014; Nakabachi 2015). Similarly, we found two large insertions of *Wolbachia* in two termite genomes. The first large insertion was in position 1,613,813 to 1,730,938 of a 1,877,474 bp scaffold of *Leptomyxotermes doriae*. No other region of the scaffold aligned to *Wolbachia* genomes. Among the alignments between this fragment and *Wolbachia* genomes, 16 were 1,981 to 20,755 bp in length, with DNA identities ranging between 31.97% and 56.53%. All these alignments were with a genome belonging to the supergroup F, which is widespread in termites (Hellemans et al. 2019; Lo and Evans 2007; Salunke et al. 2010; Yashiro and Lo 2019). The fragment contained one pseudogene of *Wolbachia* origin and 52 genes, 45 of which were confirmed to be of *Wolbachia* origin and accounted for 84.91% of horizontally acquired genes in *L. doriae* (Table S8).

The second large *Wolbachia* insertion was in a 44,091,414 bp scaffold of *Incisitermes schwarzi*, between positions 44,003,546 and 44,091,393. We generated 35 long alignments between this fragment and *Wolbachia* genomes with lengths of 2,045 bp to 15,626 bp and DNA identities of 23.91% to 59.69%. All these alignments were with strains of supergroups A and B, which have been found in various termites (Baldo et al. 2006; Lo and Evans 2007; Roy et al. 2015). This fragment contained two pseudogenes of *Wolbachia* origin and 11 genes, including ten confirmed as horizontally acquired from *Wolbachia*, which accounted for 71.43% of all horizontally acquired genes in *I. schwarzi* (Table S9).

Overall, our results showed that large insertions contributed to most horizontally acquired genes found in *L. doriae* and *I. schwarzi*. The presence of many genes and few pseudogenes in these insertions suggest a recent acquisition. These results indicate that the abundance of horizontally acquired genes in these genomes is explained by single acquisitions of large regions rather than an increased frequency of small HGTs.

### Many sub-orthogroups originated from multiple independent horizontal gene transfer events in various termite lineages

We identified 168 sub-orthogroups forming sister clades to non-metazoan sequences. For each sub-orthogroup, we estimated the number of HGT events along the species tree using the maximum-likelihood-based methods implemented in Badirate (BD-ML) and the R package ape (Ape-ML) and the parsimony-based method implemented in Badirate (BD-P). Both genic and pseudogenic copies of each sub-orthogroup were considered. BD-ML failed for one sub-orthogroup (HOG0015084.2) acquired from undetermined *Pseudomonadota* represented by highly heterogeneous copy numbers among species; however, the other two methods supported an ancient origin from an HGT event predating the divergence of termites and the two cockroach outgroups and one loss in *Marginitermes hubbardi*.

A total of 103 sub-orthogroups were species-specific, including 84 inferred to originate from a single HGT event by the three methods. These 84 sub-orthogroups had no pseudogenic copies and only one genic copy, except for two sub-orthogroups with two genic copies, suggesting recent HGT events. The remaining 19 species-specific sub-orthogroups were inferred by at least one method to originate from a single ancient HGT event retained by one species and lost by its sister group (Table S10). Eight of these sub-orthogroups were specific to *B. orientalis*, one to *M. darwiniensis*, two to *S. victoriensis*, two to *H. sjostedti*, and six to *Porotermes adamsoni*. They were all multi-copy except for two sub-orthogroups specific to *B. orientalis*. These results suggest that the inference of HGT events was generally accurate, except perhaps for multiple-copy sub-orthogroups specific to a single species lying on a long branch in the species tree.

The remaining 64 sub-orthogroups present in more than one species (with HOG0015084.2 excluded due to failure of BD-ML) were generally inferred to originate from multiple independent and species-specific HGT events (Figures 3A-C). BD-ML inferred 303 HGT events (4.73 per sub-orthogroup on average), 243 (80.20%) of which were species-specific. Ape-ML inferred 202 HGTs (3.16 per sub-orthogroup on average), including 133 (65.84%) species-specific events. BD-P inferred 228 HGT events (3.56 per sub-orthogroup on average), and 182 (79.82%) were species-specific. Notably, the prevalence of sub-orthogroups among termite and cockroach species was highly predictive of the number of HGT events inferred for each sub-orthogroup (*ρ* = 0.932 for BD-ML, *ρ* = 0.877 for Ape-ML, and *ρ* = 0.900 for BD-P, *P* < 0.001 in all cases). Overall, these results suggest that most sub-orthogroups present in multiple genomes originated from multiple independent HGTs, including many specific to a single species.

**Figure 3.**
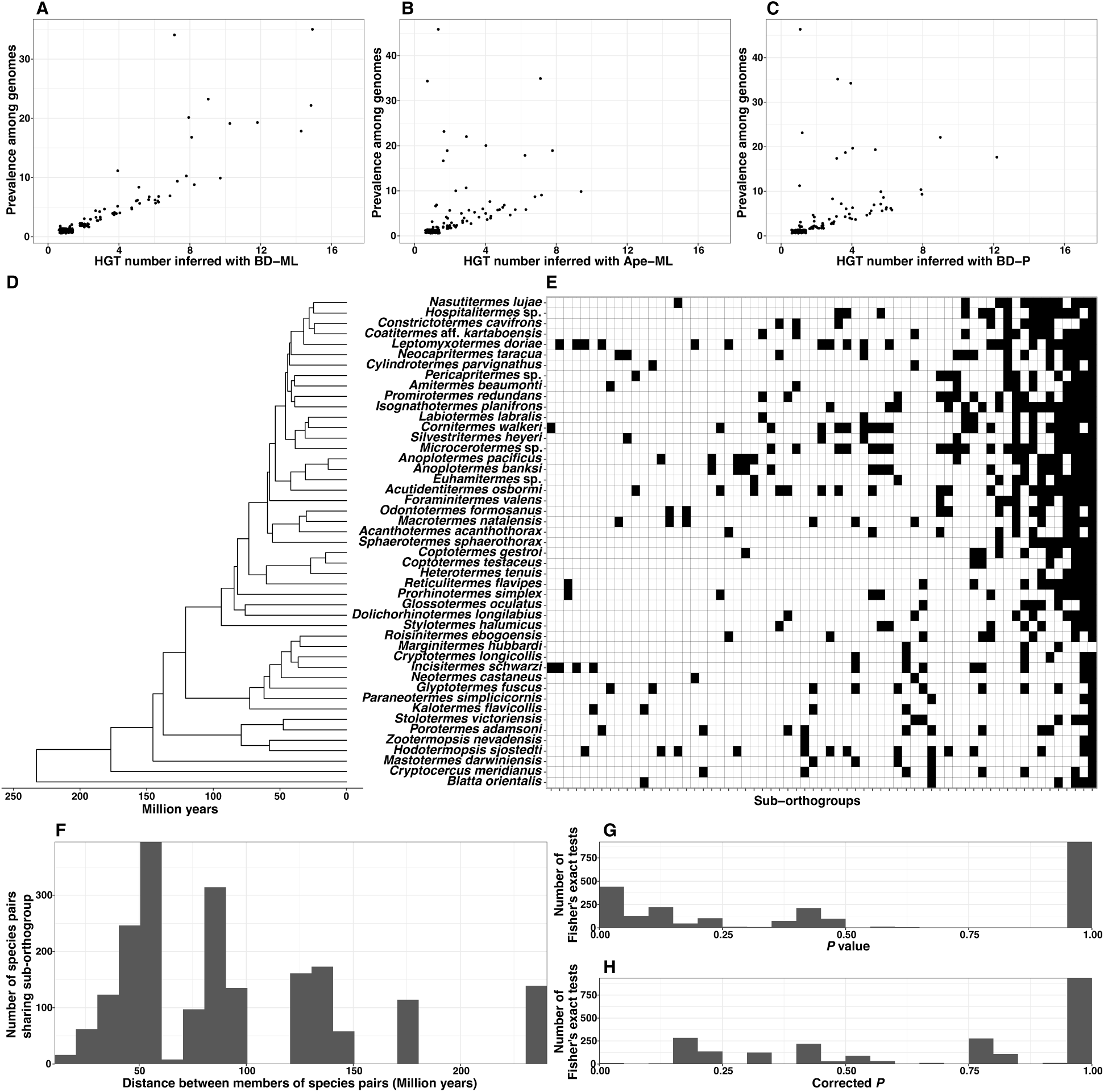
Sub-orthogroups present in multiple genomes originated via multiple independent HGTs. Relationship between the prevalence of sub-orthogroups among termites and the number of HGTs events inferred by (A) BD-ML, (B) Ape-ML, and (C) BD-P. (D) Time-calibrated species tree of 45 termites and two cockroaches. (E) Distribution of the 65 horizontally acquired sub-orthogroups found in more than one genome. Black indicates presence and white absence. (F) Distribution of divergence time between each species having a sub-orthogroup and its closest relative sharing the same sub-orthogroup. Distributions of (G) P values and (H) corrected P values of 2,251 Fisher’s exact tests testing whether sub-orthogroups were consistently associated with specific flanking HOGs.

Alternative lines of evidence supported that the sub-orthogroups present in multiple species resulted from multiple independent HGT events. First, they were generally found in distantly related species (Figures 3D-E). For example, among the 20 sub-orthogroups present in two species, 12 were shared by species that diverged over 100 million years ago (Mya) while missing in the close relatives of these species. We systematically inspected the divergence time of each species having a sub-orthogroup and its closest relative sharing the same sub-orthogroup (Figure 3F). Among the 2,041 species pairs, 645 (32.04%) had divergence times exceeding 100 Mya, and 949 species pairs (46.50%) had divergence times varying between 50 and 100 Mya. Therefore, these sub-orthogroups were generally present in unrelated termite species, indicating multiple independent HGTs rather than ancestral HGTs followed by multiple independent losses. Second, we inspected whether the location of gene and pseudogene copies of each sub-orthogroup were conserved across genomes. We tested whether sub-orthogroups were consistently associated with specific flanking HOGs using Fisher’s exact tests composed of four categories to which every 47 genomes were assigned: (1) presence of the sub-orthogroup flanked by the HOG; (2) presence of the sub-orthogroup not flanked by the HOG; (3) absence of the sub-orthogroup and presence of another sub-orthogroup flanking the HOG; (4) absence of the sub-orthogroup with no other sub-orthogroups flanking the HOG. We performed 2,288 tests, one for each observed association of a specific sub-orthogroup with a specific flanking HOG. In total, 28 tests yielded *P* values below 0.01, and 441 *P* values below 0.05 (Figure 3G). These numbers went down to 6 and 10 after correcting *P* values for multiple hypothesis testing (Figure 3H), indicating a loose association between sub-orthogroups and HOGs. Therefore, the copies belonging to the same horizontally acquired sub-orthogroups were generally flanked by different HOGs, suggesting multiple independent acquisitions by HGT in termite and cockroach genomes.

Overall, our results revealed that about two-thirds of HGT events were recent and species-specific and that sub-orthogroups found in multiple lineages often originated from independent HGT events. The species-specific nature of most genes acquired by HGT aligns with observations in other organisms. For example, almost 80% of the HGTs detected by Li et al. (2022) in 218 insect species were species-specific, and most HGTs between prokaryotes and eukaryotes are recent (Katz 2015). Rapid loss of horizontally acquired genes was also reported in bacteria and plants (Choi and Kim 2007; Liu et al. 2004; Yang et al. 2019; Dorrell et al. 2021). The rarity of functional HGTs maintained for hundreds of millions of years in large clades of eukaryotes suggests that HGTs are often lost (Husnik and McCutcheon 2018), as we also demonstrate here in termite and cockroach genomes.

### Mobile genetic elements and genetic information processing genes experience frequent horizontal transfers

Some horizontally acquired sub-orthogroups were more frequently transferred than others. We investigated whether the functions of sub-orthogroups affect their propensity for HGT. We considered 281 HGT events for the 168 sub-orthogroups (Table S11), which were determined based on manual curations of the results of BD-ML, BD-P, and Ape-ML. Our consensus set of HGTs included 190 HGTs identified by all three methods and 57 HGTs recovered by two methods. Each HGT event was associated with a branch of the time-calibrated species tree.

We first used gene functions predicted with the KEGG database. A total of 39 KEGG orthologues (KOs) were assigned to 53 sub-orthogroups, accounting for 98 HGT events. Among them, K14744 (*rzpD*; prophage endopeptidase) had the highest HGT frequency, represented by ten (18.87%) sub-orthogroups accounting for 37 (37.76%) HGT events from undetermined *Pseudomonadota*. Three transposase KOs (K07486, K07493, and K07494) were assigned to six sub-orthogroups (11.32%) and 12 HGT events (12.24%) from *Wolbachia*. Finally, 17 genetic information processing KOs, including DNA replication and repair, transcription, and translation, were associated with 18 sub-orthogroups (33.96%) and 21 HGT events (21.43%) (17 from *Wolbachia*, three from fungi, and one from *Mycoplasmatota*).

We also investigated gene functions using the Pfam database. A total of 116 Pfam domains were annotated in 121 sub-orthogroups, accounting for 214 HGT events. Five Pfam domains related to phage and prophage (PF03245, PF04860, PF05136, PF13547, and PF13550) were associated with 15 sub-orthogroups (12.40%) and 42 (19.63%) HGT events (37 from undetermined Pseudomonadota and five from *Wolbachia*). Nine Pfam domains related to transposons (PF00872, PF01348, PF01548, PF01609, PF01710, PF02371, PF12784, PF13358, and PF13655) were associated with 13 sub-orthogroups (10.74%) and 26 HGT events (12.15%) from *Wolbachia*. A total of 36 Pfam domains related to genetic information processing were assigned to 37 sub-orthogroups (30.58%) and 54 HGT events (25.23%) (26 from *Wolbachia*, 22 from viruses, four from fungi, one from plants, and one from *Mycoplasmatota*). In addition, genes encoding ankyrin repeats (PF00023, PF12796, PF13637, and PF13857) were assigned to seven sub-orthogroups (5.79%) and 16 HGT events (7.84%), including 14 from *Wolbachia*, one from undetermined *Pseudomonadota*, and one from Fungi. Therefore, mobile genetic elements and genetic information processing genes were frequently transferred to termite genomes.

The rampant recurrent transfers of mobile genetic elements and genetic information processing genes were not driven by selection for the acquisition of new genes with new functions. Indeed, most horizontally acquired genes of similar functions were present in distantly related species and absent in their respective sister groups (Figure 4), and many sub-orthogroups of HGT origin were only represented by pseudogenes, indicating that most horizontally acquired genes are transient. Instead, the frequent horizontal transfer of mobile genetic elements and genetic information processing genes may be explained by the unique biology of the major donor of HGTs to termites, *Wolbachia*. Note that the very frequent horizontal transmission of genes among bacteria themselves, especially those related with phage and prophage, often prevents more precise taxonomic assignment than *Pseudomonadota*, the bacterial phylum to which *Wolbachia* belongs (Kent and Bordenstein 2010). *Wolbachia* also undergoes rampant host-switching (Kaur et al. 2021), leading to unrelated termite species hosting closely related *Wolbachia* strains (Lo and Evans 2007), thence enabling recurrent transfers of related sequences. Furthermore, up to 20% of *Wolbachia* genomes are composed of mobile genetic elements, including prophages, transposons, and group II introns, and ankyrin repeats account for up to 4% of protein-coding genes in *Wolbachia* genomes (Wu et al. 2004; Iturbe-Ormaetxe et al. 2005; Klasson et al. 2008, 2009; Kent and Bordenstein 2010; Cerveau et al. 2011; Leclercq et al. 2011; Siozios et al. 2013; Kaur et al. 2021). The abundance and mobile nature of these genes could explain their frequent insertions inside termite genomes. Finally, the frequent HGTs of genes related to genetic information processing and prophage genes, especially those related to host cell lysis, might be related to the phage-mediated lysis of *Wolbachia*. These genes are expected to be highly expressed during the lytic stage, during which *Wolbachia* cells break and release numerous phage particles and *Wolbachia* DNA into host cells, increasing the probability of HGTs. Overall, our results highlight that the frequent transfers of mobile genetic elements and genetic information processing genes to the host genome are not functional.

**Figure 4.**
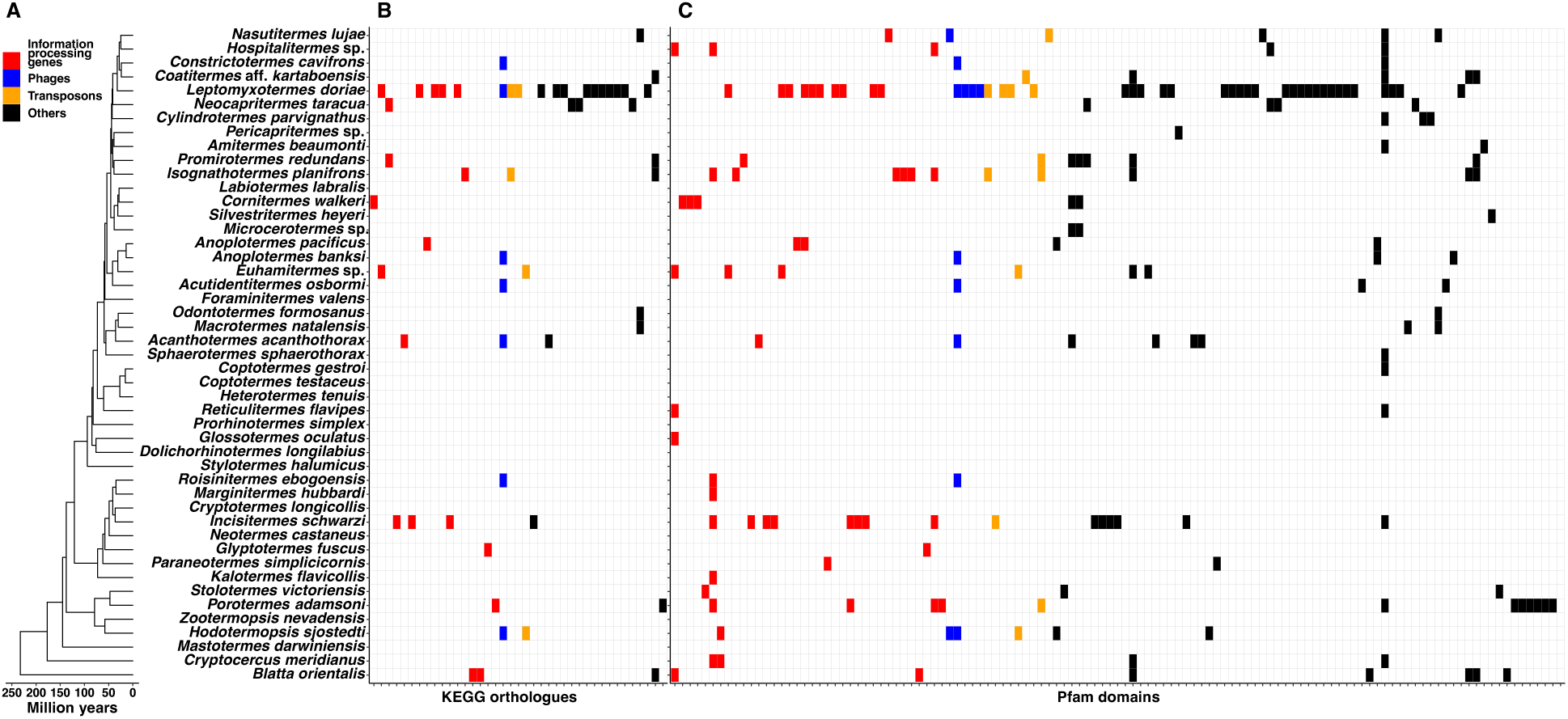
Distribution of horizontally acquired genes with functions annotated using the KEGG and Pfam databases. (A) Time-calibrated species tree of 45 termites and two cockroaches. Filled and empty cells indicate the presence and absence of horizontally acquired genes associated with (B) KEGG orthologues and (C) Pfam domains. Red cells, information processing genes; blue cells, phage; yellow cells, transposons; black cells, others.

### Horizontally acquired genes experience frequent duplications and pseudogenizations

We investigated the evolution of horizontally acquired sub-orthogroups using the midpoints of the branches to which HGT events were associated in the time-calibrated species tree as estimations of HGT times. Out of the 591 records of the 168 sub-orthogroups among our 47 genomes, 240 (40.61%) were represented by more than one copy and 29 (4.91%) by more than 20 copies (Figure 5A). 383 records (64.81%) were only represented by pseudogenic copies and 168 (28.43%) only by genic copies (Figure 5B). Both the copy number of each sub-orthogroup per genome and the proportion of pseudogenes were positively correlated with HGT times (*P* < 0.001) with *ρ* values of 0.510 and 0.553, respectively. These results showed that duplications and pseudogenizations are common in horizontally acquired genes, as previously observed in various organisms (Nikoh et al. 2010; Paganini et al. 2012; Sun et al. 2013a, 2013b; Klasson et al. 2014; Savory et al. 2015; Husnik and McCutcheon 2018; Dai et al. 2021). The extensive pseudogenization observed here is in line with the high turnover of most horizontally acquired genes amongst termite species.

**Figure 5.**
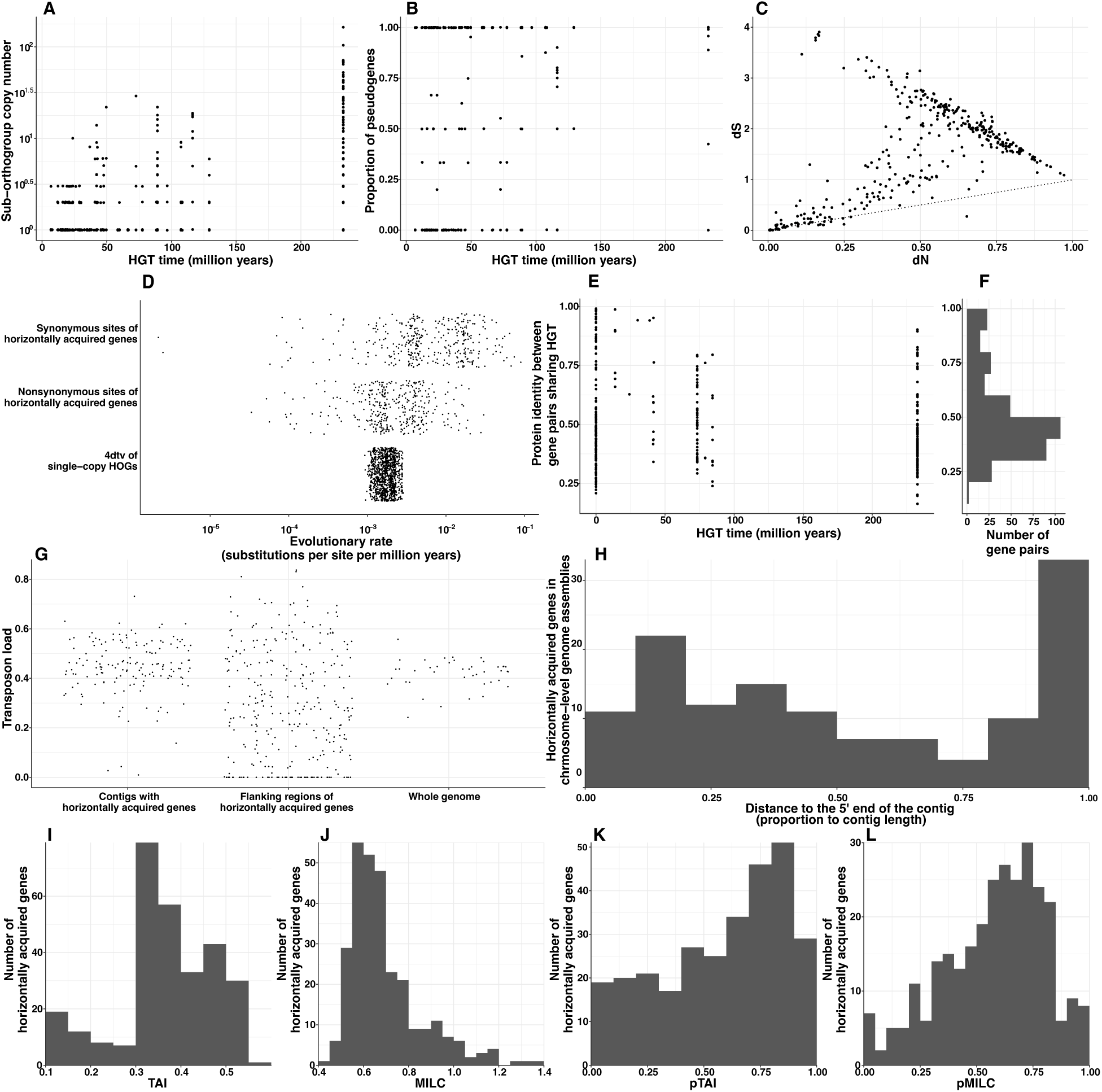
Evolution of horizontally acquired genes after HGT events. (A) Copy number and (B) proportion of pseudogenes in each genome for each sub-orthogroup acquired by HGT plotted against the time of HGT events. (C) Synonymous and nonsynonymous substitution rates (*dS* and *dN*) for gene pairs descending from single HGT events (the dot line indicates dN/dS = 1). (D) Evolutionary rates of synonymous and nonsynonymous sites of horizontally acquired genes and four-fold degenerated synonymous sites (4dtv) of single-copy HOGs shared by the 47 genomes. Protein identity of gene pairs descending from single HGT events plotted against (E) the time of HGT events and (F) the distribution of protein identity. (G)Location of horizontally acquired genes in the 22 chromosome-level genome assemblies. (H) Transposon load of the contigs containing horizontally acquired genes, the 5-kb upstream and downstream regions flanking horizontally acquired genes, and the whole genome. Distribution of (I) the tRNA adaptation index (TAI), (J) the measure independent of length and composition (MILC), (K) the percentile TAI (pTAI), and (L) the percentile MILC (pMILC) of horizontally acquired genes.

### Horizontally acquired genes evolve faster than synonymous sites of single-copy protein-coding genes

We estimated synonymous and nonsynonymous substitution rates (*dS* and *dN*) between pairs of genes descending from single HGT events to investigate the selection pressures operating on horizontally acquired genes. We used 360 gene pairs, 339 (94.17%) of which had *dN*⁄*dS* values below 1 (Figure 5C), suggesting, at first glance, purifying selection on most horizontally acquired genes. However, the evolutionary rate of synonymous and nonsynonymous sites of horizontally acquired genes were both significantly higher than that of the four-fold degenerated sites (4dtv) of single-copy HOGs (Wilcoxon test, *P* < 0.001) (Figure 5D). Furthermore, the protein identity between genes forming a pair was low, ranging between 16.35% and 99.11% (mean: 50.66%, median: 45.90%), and was weakly correlated with HGT time (*ρ* = −0.131, *P* = 0.013) (Figures 5E-F). Therefore, the three codon positions of horizontally acquired gene sequences evolved faster than synonymous sites of native protein-coding genes, suggesting horizontally acquired genes are not under purifying selection. *dN*⁄*dS* values lower than 1 have been reported for horizontally acquired genes using both phylogenomic or population genetic methods in various eukaryotic lineages, such as whiteflies (Gilbert and Maumus 2022), Lepidoptera (Sun et al. 2013b), fungi (Sun et al. 2013a), diatoms (Vancaester et al. 2020), and plants (Guo et al. 2023). They have often been interpreted as evidence of purifying selection on horizontally acquired genes, suggesting the functionalization of these genes. However, our estimations of evolutionary rates in absolute time show that both synonymous and nonsynonymous sites of horizontally acquired genes accumulate substitutions at a faster rate than four-fold degenerated sites of native protein-coding genes, indicating they are not under purifying selection but instead experience accelerated evolution.

Repeat-rich genomic regions often evolve rapidly (Henikoff et al. 2001; Bosco et al. 2007; Brand and Levine 2021), which could provide an explanation for the high evolutionary rates of horizontally acquired genes. However, we found that the transposon load of the 5-kb upstream and downstream regions flanking horizontally acquired genes was significantly lower than the transposon loads of the contigs and the whole genome (paired Wilcoxon test, *P* < 0.001) (Figure 5G). We also inspected the 22 chromosome-level genome assemblies and found that horizontally acquired genes were located along the entire contigs (Figure 5H), without clear patterns of association with the centromere or telomere regions. Therefore, the position of horizontally acquired genes in the genome is unlikely to explain their accelerated evolution, which should result from a yet unknown mechanism.

### Horizontally acquired genes do not undergo post-HGT codon usage optimization

We found that the synonymous sites of horizontally acquired genes evolved faster than 4dtv of single-copy HOGs, which may promote codon usage optimization —codon usage approaching that of termites and cockroaches as substitutions are accumulated. We inspected codon usage of horizontally acquired genes using the tRNA adaptation index (TAI) (Reis et al. 2004) and the measure independent of length and composition (MILC) (Supek and Vlahoviček 2005). High TAI and low MILC reflect codon usage optimization. In addition, for each horizontally acquired gene, we also calculated the proportion of genes having a lower TAI (pTAI) and MILC (pMILC) in the same genome. TAI values of horizontally acquired genes ranged between 0.117 and 0.554 (Figure 5I), and MILC values ranged between 0.417 and 1.367 (Figure 5J). pTAI values of horizontally acquired genes ranged between 0.008 and 0.986 (Figure 5K), and pMILC values ranged between 0.002 and 0.991 (Figure 5L), suggesting that their codon usage are not distinguishable from that of other termite and cockroach genes. In addition, the correlations between the time of HGT and codon usage indices provided conflicting results. HGT time positively correlated with TAI (*P* < 0.001, *ρ* = 0.608), did not correlate with MILC (*P* = 0.06, *ρ* = 0.111), negatively correlated with pTAI, albeit weakly (*P* = 0.002, *ρ* = −0.185), and presented a significant but weak positive correlation with pMILC (*P* = 0.011, *ρ* = 0.150). While some horizontally acquired genes may occasionally present optimized codon usage, most show no clear evidence of post-HGT optimization. The codon usages of horizontally acquired genes spanned the entire spectrum of codon usages found in native termite and cockroach genes, providing an explanation for the failure of past attempts to detect HGT based on codon frequency (Friedman and Ely 2012).

### Introns are both transferred with horizontally acquired genes and acquired after HGT

We inspected horizontally acquired genes for the presence of introns. We found a total of 173 introns in 88 horizontally acquired genes. Among them, 80 genes possessed one to three introns, and eight possessed more than three introns. We searched all introns against NCBI’s nt database and found that 26 introns only yielded homology hits with insects, one intron yielded 48 homology hits with insects and one with ostracods, and one intron yielded 84 homology hits with arthropods, three hits with *Wolbachia*, and three hits with viruses. These introns were probably native to insect genomes and gained after HGT events. They belonged to 22 genes found across unrelated genera, including *Blatta*, *Hodotermopsis*, *Paraneotermes*, *Anoplotermes*, *Isognathotermes*, *Amitermes*, *Coatitermes*, and *Nasutitermes*. Among these genes, there was one transcription factor of fungal origin in *B. orientalis* (*Bori00029538-R0*, KO: K09228, K09191; Pfam: PF00096), two thaumatin genes of plant origin (Pfam: PF00314) in *A. pacificus* (*Apac00010071-R0*) and *A. banksi* (*Aban00013346-R0*), one tRNA methyl transferase HUP domain originating from *Wolbachia* (Pfam: PF03054) in *Nasutitermes lujae* (*Nluj00017428-R0*) and six ankyrin repeat genes, two of bacterial origin in *Coatitermes* aff. *kartaboensis* (*Csp400006450-R0*) and *Isognathotermes planifrons* (*Iunk00007542-R0*) and four of fungal origin in *B. orientalis* (*Bori00027419-R0*, *Bori00027420-R0*, *Bori00027340-R0*, and *Bori00011858-R1*). Another 27 introns were associated with *Wolbachia*, including 13 introns with homology hits with *Wolbachia* only and 14 introns with more than 60% of homology hits with *Wolbachia*. These introns were presumably acquired together with the genes within which they reside during HGT.

## Conclusions

Our study sheds light on the origin and prevalence of HGTs from non-metazoan organisms to termite and cockroach genomes as well as the fate of these horizontally acquired genes. We found that, like all organisms, termites and cockroaches have acquired genes by HGT during their evolutionary history. However, several lines of evidence indicate that most genes acquired horizontally by termites and cockroaches are non-functional: (1) the analyses of 47 genomes revealed 271 genes and 2,479 pseudogenes originating from HGTs, suggesting high pseudogenization rates following HGT; (2) many pseudogenes did not have genic copies in the same genome; (3) horizontally acquired genes, including orthologous genes, generally originated from independent HGTs, unique to each termite species, suggesting recent acquisitions; (4) the substitution rate of horizontally acquired genes was faster than synonymous sites of native protein-coding genes; and (5) the protein sequences encoded by genes descended from the same HGT events were not conserved. These results indicate that most horizontally acquired genes are non-functional, likely to be lost, and have therefore not played an important role in termite evolution. Instead, horizontally acquired genes are often mobile genetic elements, genes involved in genetic information processing, and ankyrin repeats, whose frequent transmission is dictated by non-selective processes, such as high expression during the lytic stage of *Wolbachia*.

## Materials and methods

### Genome data

We used the genome assemblies and annotations of 45 termites and two cockroaches and the corresponding time-calibrated phylogenetic tree generated by Liu et al. (2025). The tree was used to cluster protein-coding genes of all genomes into hierarchical orthologous groups (HOGs) using OrthoFinder v.2.5.4 (Emms and Kelly 2019). Gene functions were inferred using InterproScan (Jones et al. 2014) and KOfamScan (Aramaki et al. 2020). InterproScan annotates protein domains based on the Pfam database (Bateman et al. 2004), while KOfamScan provides annotations based on KEGG Orthology (Kanehisa 2002). Transfer RNA (tRNA) genes were annotated using tRNAScan-SE v.2.0.11 (Chan et al. 2021) with the eukaryotic model and filtered with EukHighConfidenceFilter implemented in tRNAScan-SE. Transposons were annotated using the sensitive mode of EDTA v.2.2.0 (Ou et al. 2019).

### Identification of horizontal gene transfers

We screened the genomes of 45 termites and two cockroaches for horizontal gene transfers (HGTs) from organisms other than Metazoa. Potential HGTs were initially identified using (1) the alien index (AI), which compares the *e*-values of the best non-arthropod metazoan homology hits to the best non-metazoan hits, and (2) the proportion of homology hits from non-arthropod metazoan and non-metazoan organisms. We searched the entire gene repertoire of each genome against the non-redundant (nr) protein database (accessed in July 2022) using DIAMOND v2.1.7.161 with the arguments “*–max-target-seqs 500 –min-score 50*” (Buchfink et al. 2015). The AI of each query sequence was calculated with the *e*-value of the best hit to non-arthropod Metazoa (*bh_in_*) and the *e*-value of the best hit to organisms other than Metazoa (*bh_out_*), using the equation (Gladyshev et al. 2008; Yuan et al. 2023):

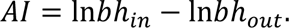

Sequences were retained as HGT candidates if (1) their AI was above 45, (2) they had at least 25 homology hits, and (3) more than 90% of the homology hits were with non-metazoan sequences (Gladyshev et al. 2008; Yuan et al. 2023).

To validate HGT candidates and identify putative donor taxa, we performed phylogenetic analyses of the HOGs containing HGT candidates together with homologous sequences obtained from the nr database. HOGs containing at least one HGT candidate were searched against the nr database using DIAMOND with the arguments “*–max-target-seqs 100 –min-score 50*”. In the case of homology hits from various sequences of the same species, only the sequence that appeared first in the DIAMOND output was retained, and the remaining sequences were dropped since they increased the computational burden without providing extra information on donor taxa. Sequences identified with DIAMOND were removed if they contained ambiguous amino acids (B for asparagine/aspartic acid and Z for glutamine/glutamic acid) or were labelled as “partial” in the titles. The protein sequences of each HOG and their homologous sequences identified with DIAMOND were aligned using MAFFT v.7.508 (Katoh et al. 2002) with the *–auto* option. Protein sequence alignments were trimmed using trimAL v.1.4.rev22 with the *-automated1* option (Capella-Gutiérrez et al. 2009), which removes spurious sequences and regions. Phylogenetic trees were inferred from the trimmed protein alignments using IQ-TREE v.2.2.0.3 (Minh et al. 2020) with the best-fitting amino acid substitution models for nuclear genomes selected by ModelFinder (Kalyaanamoorthy et al. 2017) and 1,000 bootstrap replicates. The phylogenetic trees were rooted at the midpoint using the *midpoint.root* function implemented in R package *phytools* (Revell 2024) and manually inspected in order to validate HGTs and identify putative donor taxa. HOGs were considered horizontally acquired when all their sequences were nested within non-metazoan sequences. We further divided HOGs into sub-orthogroups when termite sequences formed multiple clades separated by non-metazoan sequences.

We identified pseudogenetic copies of horizontally acquired genes using PseudoPipe (Zhang et al. 2006). Briefly, the protein sequences of all horizontally acquired genes identified in this study were aligned to all 47 genomes with tBLASTn (Altschul et al. 1997). This enabled the identification of unitary pseudogenes having no gene orthologue in the given genome. BLAST hits with scores above 50, *e*-value below 1*e* − 10, and query coverage above 75% were processed with PseudoPipe, which provides complete annotation of the pseudogenes, including their locations in the genome and the names of their parental genes. Pseudogenes that matched with less than 50% of the parental genes in terms of protein length were not considered.

### Validation of HAGPs

Some HGT candidates identified above may be foreign DNA sequences that have contaminated the genome assemblies. Therefore, the HGT candidates were further validated as follows. We inspected the contigs containing horizontally acquired gene and pseudogene candidates to confirm they are of termite and cockroach nuclear genome origin. We used five criteria to filter putative contaminations and validate HGT candidates. Contigs with HAGPs were retained if they met the following criteria: (1) longer than 25 kb; (2) coding density estimated with PRODIGAL v.2.6.3 (Hyatt et al. 2010) lower than 0.7; (3) horizontally acquired genes making up less than 70% of the contig genes; (4) containing at least two genes not horizontally acquired; and (5) having a ratio of contig sequencing depth (the ratio between the total length of reads mapping to the contig and the length of the contig) to genome sequencing depth above 0.5 and below 10 in a set of deep long sequencing from Liu et al. (2025) (Table S1). We estimated the genome and contig sequencing depth with SAMtools v.1.12 (Li et al. 2009) using long reads (Table S1) mapped to the corresponding genomes with minimap2 2.24 (Li 2018). In addition to the five criteria described above and used for both contig filtering and validation, we inspected a series of other parameters to ensure the reliability of our detection procedure. We inspected the coding density, horizontally acquired gene length, and proportion of transposons as additional evidence of the accuracy of our HGT detection procedure. We also searched the genome assemblies against genomes of *Wolbachia* (Table S12), termite gut bacteria (Table S5), and *Blattabacterium* (Table S6) using minimap2, and calculated the proportion of the contigs aligning to these bacterial genomes. All reference bacterial genomes were downloaded from NCBI. The *Wolbachia* genomes belonged to 17 strains from the supergroups A-F, which are representative of the *Wolbachia* diversity (Table S12). The termite gut bacterial genomes included chromosome-level assemblies of one *Breznakiella*, one *Fibrobacter*, one *Gracilinema*, and ten *Treponema* species (Table S5). The *Blattabacterium* genomes included one strain from *Mastotermes*, six from *Cryptocercus*, one from *Blatta orientalis*, and nine from other cockroaches (Table S6). Finally, we searched for the presence of HGTs in conspecific samples by mapping long and short reads generated by Liu et al. (2025) (Table S13) using minimap2. The sequencing depth and coverage (proportion of a region with sequencing depth above 0) were calculated for the contigs and the horizontally acquired genes/pseudogenes using SAMtools.

### Modelling horizontal gene transfer events

Some horizontally acquired HOGs were composed of several sub-orthogroups, suggesting their parallel origin from independent HGTs. Similarly, some sub-orthogroups may be the results of multiple HGTs from closely related donors rather than a single HGT. We investigated the dynamic of HGTs along the species tree using the total copy number (genes and pseudogenes) found in the genomes of extant species. The analyses were performed for each sub-orthogroup separately using three methods to estimate the number of HGT events they each experienced. We performed ancestral state reconstruction using the *ace* function implemented in the R package *ape* (Paradis and Schliep 2019) with a Brownian motion model fitted by the maximum likelihood (ML) method. We also estimated HGT turnover rates with Badirate v.1.35.00 (Librado et al. 2012), using the ML-fitted birth, death, and innovation (BDI) model with the free rate (FR) branch model, which assumes different BDI rates among species tree branches. Finally, we used the parsimony-based method implemented in Badirate.

### Analyses of selection, codon usage, and introns of horizontally acquired genes

We investigated the evolution of horizontally acquired genes following their insertions in termite and cockroach genomes. Three aspects of horizontally acquired gene evolution were studied: (1) the evolution of protein-coding sequences, (2) the evolution of codon usage, and (3) the acquisition of introns.

For (1), we compared protein-coding sequences that descended from a single HGT event to ensure the measured signal represented the evolution of sequences residing inside termite genomes. To do so, we manually curated HGT events inferred by Badirate and the R package *ape* and split sub-orthogroups into groups of genes and pseudogenes that originated from a single HGT event. We computed pairwise protein alignments for these groups of genes and pseudogenes using MAFFT with the parameter “*–auto*” and converted the protein alignments into codon alignments with PAL2NAL v.14 (Suyama et al. 2006). The codon alignments were used to estimate synonymous and nonsynonymous substitution rates (*dS* and *dN*) using KaKs_calculator 3.0 with default settings (Zhang 2022), and the protein alignments were used to compute protein identity using the *dist.alignment* function with *matrix=“identity”* implemented in the R package *seqinr* (Charif and Lobry 2007). We compared the substitution rate of horizontally acquired genes to the neutral substitution rate of native termite and cockroach genes. The native neutral substitution rates were estimated using single-copy HOGs. Briefly, we aligned protein sequences of 1,410 single-copy HOGs using MAFFT with the parameter “*–auto*”, converted the protein alignments into codon alignments with PAL2NAL, concatenated the codon alignments into a supermatrix, and extracted a total of 41,729 four-fold degenerated synonymous sites (4dtv). We used the species tree generated by Liu et al. (2025) and the alignment of 4dtv to estimate branch length using IQ-TREE with the setting “*-mset GTR”*, which limits the nucleotide substitution model selection by ModelFinder to those of the general time reversible (GTR) family. ModelFinder selected the GTR+F+R4 model. The neutral substitution rate of species pairs was estimated as the halved ratio of pairwise divergence in the non-ultrametric 4dtv tree and the time-calibrated tree.

For the analysis of codon usage (2), we computed two indices. The first index was the tRNA adaptation index (TAI) computed using the R package *tAI* (Reis et al. 2004). TAI compares the codon usage of a gene to the genomic tRNA copy number. A high TAI value is indicative of a gene highly co-adapted to the tRNA pool (Reis et al. 2004). The second index was the Measure Independent of Length and Composition (MILC) (Supek and Vlahoviček 2005) calculated using the R package *coRdon* (Elek et al. 2018). MILC quantifies the dissimilarity between the codon usage of a target gene and the ribosomal genes, which are highly expressed housekeeping genes expected to display near-optimal codon usage. A low MILC value indicates a high similarity to the codon usage of ribosomal genes (Supek and Vlahoviček 2005). We identified termite and cockroach ribosomal genes by searching the entire gene repertoire of each genome against the nr (accessed in July 2022) database using DIAMOND with the arguments “*--min-score 50 --query-cover 75”* and extracted the best homology hits containing the term “ribosomal protein” in the titles.

For the analysis of introns (3), we extracted the intron sequences of horizontally acquired genes and carried out searches against the NCBI nucleotide (nt) database (accessed in July 2022) using BLAST 2.13.0+ (Camacho et al. 2009) with the arguments “*-evalue 1e-10 -max_target_seqs 25”*.

### Statistical analyses

All statistical analyses were performed with R v.4.2.1 (R Core Team 2013). Spearman’s rank correlation coefficients and their significance were computed using the R function *cor.test*. Fisher’s exact tests were performed with the R function *fisher.test*. For multiple hypothesis testing, the false discovery rate (FDR) was computed using the R function *p.adjust* with the option *method=“BH”*. Wilcoxon tests were performed using the R function *wilcox.test* with the argument “*paired=TRUE”*. Linear models were fitted using the R function *lm*.

## Acknowledgements

We thank OIST’s Scientific Computation and Data Analysis Section (SCDA) for providing access to the OIST computing cluster. This work was supported by subsidiary funding from OIST, including funding for a workshop held at OIST in early December 2022 and funding by the Deutsche Forschungsgemeinschaft (DFG, German Research Foundation) to DPM (MC 436/5-1 and MC 436/7-1) and MCH (HA 8997/1-1). This work was also supported by the Czech Science Foundation (project No. 24-212674S to T.B.).

## Author Contributions Statement

CL, SH, YK, MCH, DPM, and TB conceptualised the experiments. CL analysed the data. CL and TB wrote the original draft manuscript. SH, YK, AAM, CA, YW, AB, FH, JŠ, MCH, and DPM revised the manuscript. All authors (CL, SH, YK, AAM, CA, YW, AB, FH, JŠ, MCH, DPM, and TB) read and accepted the final version of this manuscript.

